# Development of Specific barcodes for identification of *Aloe species* based on chloroplast DNA barcoding

**DOI:** 10.1101/2023.07.24.550245

**Authors:** Subrata K. Das, Alpana Joshi

## Abstract

DNA barcoding is currently an effective and extensively used tool for species identification. The chloroplast matK (maturase K) and rbcL (Ribulose-bisphosphate large subunit) are one of the most variable coding genes of angiosperms and have been recommended as a universal barcode for land plants. *Aloe* is a large genus of around 500 species, and most species are widely used for traditional medicinal purposes worldwide, *viz*., *Aloe vera, Aloe ferox, Aloe arborescens*, and *Aloe maculate*. This study evaluated the two chloroplast DNA barcodes (matK and rbcL) to develop a theoretical base for species identification and germplasm conservation of *Aloe* species. The Maximum Parsimony analysis was conducted to study the evolutionary relatedness of *Aloe* sequences using matK and rbcL sequences retrieved from the NCBI database. The results revealed that 49 *Aloe* accessions were segregated into four major clades based on the matK sequence. Likewise, the 29 *Aloe* accessions were distributed into two major clades based on the rbcL sequence. SNP (Single Nucleotide Polymorphism) site analysis was conducted to obtain the specific barcode of *Aloe* species and generate the corresponding DNA QR code that electronic devices could immediately recognize. This study provides innovative research methods for efficient species identification of the genus *Aloe* and indicates the possibility of correctly identifying, discriminating, and documenting the *Aloe* species.

## INTRODUCTION

The *Aloe* genus is a group of shrubby succulent plants belonging to the Asphodelaceae family. *Aloe* is the largest genus, contains more than 500 species, primarily inhabitants of arid climates, and is widely distributed in Africa, India, and other Arid areas. The most significant *Aloe* species are approximately 140, and most are found in South Africa **(Grace et al., 2015)**. The most widely distributed *Aloe* species in South Africa are *Aloe greatheadii* and *Aloe ferox* **(O’Brien et al., 2011)**. *Aloe* species, namely *Aloe arborescens, Aloe barbadensis, Aloe ferox, and Aloe vera*, are commonly used as oral therapeutic agents due to their medicinal and skin care properties **(Hossain et al., 2013; Salehi et al., 2018)**. Expert botanists or analytical techniques classify most medicinal plants based on morphological characteristics. *Aloe* leaves are difficult to categorize using morphological and traditional DNA barcodes, making *Aloe* conservation and trade regulation problematic. Moreover, morphological features cannot identify the raw materials, mainly in powder or processed products obtained from plants used in herbal formulations. Spurious materials and species adulteration have become essential concerns for health and safety reasons **(Kress et al., 2015; Zhu et al., 2022; Safhi et al., 2023)**.

DNA barcoding has evolved as an efficient method for identifying medicinal plant species using short sequences. DNA barcoding enables more accurate species identification, ensuring that the products used for therapeutic purposes are authentic **(Nazar et al., 2022)**. Since the development of the plant DNA barcoding method, conserved DNA sections have been discovered in several plant species by DNA analysis, leading to the discovery of a universal DNA barcode **(Li et al., 2021; Jiang et al., 2022; Kushwaha et al., 2023)**. Consortium for the Barcode of Life (CBOL) recommended rbcL and matK regions as a standard two-locus barcode for global plant databases after evaluating the performance of seven candidate barcoding regions, namely atpF-atpH, matK, rbcL, rpoB, rpoC1, psbK-psbI, and trnH-psbA because of their ability to discriminate between species with high accuracy **(CBOL, 2009; Heckenhauer et al., 2016; Ho et al., 2021)**.

Molecular identification of six species of the genus *Aloe* was conducted based on ITS2, psbA-trnH, rbcL, and matK barcode regions **(Han, 2016)**. *Aloe saudiarabica* and *Aloe pseudorubroviolacea* are rare and endangered plant species in Saudi Arabia. All species of *Aloe*, except *Aloe vera*, are regulated by the Convention on International Trade of Endangered Species (CITES) **(Sajeva et al., 2007)**. The partial matK and rbcL gene sequence was used to discriminate and authenticate *Aloe pseudorubroviolacea* from the closely related plant species *A. rubroviolacea* **(Alaklabi et al., 2021)**. A similar study reported the identification of *Aloe saudiarabica* and *Aloe shadensis* based on plastid matK, rbcL, and the nuclear ITS region **(Algarni, 2022a; b)**. These two gene regions have also been used intensively to study the different species of the genus *Aloe* **(Grace et al., 2015; Han, 2016; Alaklabi et al., 2021; Algarni et al., 2022a; b)**. Although DNA barcoding has been studied in species identification and phylogeny construction of a few *Aloe* species, it has not been reported that DNA barcoding genes can be used to develop specific identification segments (Species-specific barcodes). The present study used matK and rbcL barcode regions to produce unique identification fragments of a particular species of *Aloe* based on phylogenetic analyses and SNP site analyses. The barcode genes were also comprehensively analyzed to obtain standard DNA marker fragments laying the foundation for evaluating, conserving, and protecting *Aloe* species.

## MATERIAL AND METHODS

### Nucleotide sequences and sequence alignment

We retrieved the matK and rbcL sequences from the NCBI Gene database (https://www.ncbi.nlm.nih.gov/) for species identification. All retrieved sequences were evaluated critically, and any low-quality sequences were removed **(Suesatpanit et al., 2017; Joshi et al., 2017)**. For analyses, sequences of each species were saved in FASTA format. The accessions numbers used for rbcL are NC_035506.1, AY323643.1, AJ512308.1, AY323647.1, AY323641.1, AJ512307.1, NC_066013.1, AY323642.1, AJ512304.1, Z73690.1, AY323640.1, AJ512306.1, AJ512283.1, AJ512302.1, AY323639.1, AJ512313.1, AY323638.1, AY323646.1, NC_044761.1, NC_035505.1, AJ512311.1, AJ512303.1, AJ512293.1, AJ512312.1, Z73680.1, AJ512288.1, AJ512292.1, Z73683.1, AJ512287.1, and AY323648.1.

The accessions numbers used for matK are NC_035506.1, KY556640.1,ON641362.1, ON641334.1, JQ276402.1, AY323726.1, AJ511390.1, GQ434051.1, KP072727.1, MW176075.1, KP072725.1, GQ434050.1, AJ511371.1, NC_066013.1, AY323723.1, AJ511412.1, AJ511425.1, KX270418.1, KX270416.1, JX903606.1, AJ511411.1, OM662302.1, AJ511385.1, AJ511389.1, AJ511387.1, AY323718.1, AY323717.1, JQ435526.1, KU748259.1, KU748254.1, AJ511388.1, KU556663.1, AJ511407.1, AY323713.1, AJ511394.1, AJ511379.1, AJ511383.1, AJ511384.1, AJ511391.1, AJ511397.1, NC_044761.1, AJ511369.1, AJ511370.1, AJ511386.1, AJ511392.1, AJ511395.1, AJ511382.1, AJ511410.1, AJ511413.1, AJ511409.1, AJ511393.1, AJ511399.1, AJ511402.1, AJ511401.1, AJ511375.1, AJ511403.1, JQ276404.1, AJ511373.1, AJ511406.1, AY323712.1, AJ511404.1, AJ511408.1, AJ511377.1, AY323704.1, AJ511376.1, AJ511380.1, AJ511372.1, AJ511381.1, KX270415.1, JQ435530.1, AY323714.1, AY323694.1, AY323695.1, AY323693.1, AY323697.1, AY323705.1, AY323706.1, AY323707.1, AY323708.1, AY323710.1, AY323711.1, AY323715.1, AY323716.1, AY323719.1, AY323720.1, AY323721.1, AY323724.1, AY323725.1, JX903603.1. NC_035505.1, JX903607.1, KC893716.1, KU748261.1, MH043303.1, KU748253.1, KU748252.1, KU748291.1, KX270417.1, AY323709.1, and KU937275.1.

### Data analysis

The sequence alignment was initially performed using the MUSCLE program of MEGA v11 with the default alignment parameters for multiple sequence alignment parameters **(Tamura et al., 2021)**. In the pairwise distances analyses, the positions containing gaps and missing were excluded from the data set (complete deletion option). Phylogenetic trees were constructed with the Maximum Parsimony method. Evolutionary divergence for each data set and pattern of nucleotide substitution was performed by MEGA 11 **(Tamura, 1992)**. The clade reliability in these trees using the Maximum Parsimony method was tested with bootstrapping with 10,000 replicates to obtain the support values of the clade nodes.

The number of INDELs for each dataset was identified by deletion/insertion polymorphisms (DIP) analysis in DnaSP v6.12 **(Rozas et al., 2017)**. The polymorphic site, genetic diversity indices, and neutrality tests, Fu & Li’s D, Fu & Li’s F, and Tajima’s D, were performed using the DnaSP v6.12 **(Li et al., 2021)**. Automatic Barcode Gap Discovery (ABGD) was conducted using ABGD webserver (URL: https://bioinfo.mnhn.fr/abi/public/abgd/abgdweb.html accessed on June 19, 2023) with default parameters using Kimura 2-Parameter (K2P) model **(Kimura, 1980; Puillandre et al., 2012)**.

## RESULTS AND DISCUSSION

### Sequence analysis

This study retrieved 49 matK sequences and 29 rbcL sequences from the NCBI Nucleotide database for further analysis. After blasting and editing, the consensus length of matK and rbcL were 835 bp and 1265 bp, respectively. The distribution of the four bases in candidate nucleotide sequences and the overall mean nucleotide base frequencies were observed for candidate nucleotide sequences using MEGA v11. The average GC contents of matK ranged from 23.08 % to 38.4 %, whereas the GC contents of rbcL ranged from 27.89 % to 58.77 % (Table 1).

**Table 1.** The nucleotide base frequencies (%) of candidate nucleotide sequences of *Aloe sp*.

| Sequences | Alignment length (nt) | Base content |  |  |  |  |  |  |  |
| --- | --- | --- | --- | --- | --- | --- | --- | --- | --- |
|  |  | T | C | A | G | GC | GC-1 | GC-2 | GC-3 |
| matK | 835 | 39.27 | 17.36 | 29.73 | 13.62 | 30.98 | 23.08 | 38.4 | 31.49 |
| rbcL | 1265 | 28.91 | 19.47 | 27.49 | 24.12 | 43.59 | 58.77 | 44.08 | 27.89 |

Polymorphism site analysis of matK and rbcL was conducted using DnaSP v6.12. The matK sequence had the highest proportion of variable sites and the lowest proportion of conservative sites as compared to the rbcL sequence **(Jiang et al., 2022; Kushwaha et al., 2023)**. The results showed 66 variable sites, 27 parsimony-informative sites, and 39 singleton sites in matK (Table 2). The conserved regions were identified in matK alignment; Region 1: 267-314 (ANTTANGNTTAACANCTTCNNCAACTTTTCTTGAACGANCCCATTTCT) and Region2 spanned from 789-835 (ATATTATTGATCGATTTGGTCGGATATGTAGAAATCTTTCTCANTAT). The 39 singleton variable sites were found at position 22, 67, 82, 86, 101, 140, 153, 175, 193, 197, 211, 266, 315, 365, 367, 404, 421, 424, 428, 439, 447, 486, 494, 545, 557, 618, 622, 664, 676, 683, 706, 745, 764, 765, 766, 767, 768, 784 and, 27 parsimony informative sites at position 50, 80, 104, 165, 167, 178, 179, 198, 199, 241, 242, 329, 387, 443, 519, 550, 590, 606, 607, 615, 620, 655, 662, 707, 715, 778, 788.

**Table 2.** The analysis of variation of candidate barcode sequences in *Aloe sp*.

| Sequence | n | Conserved site | Variable site | Parsimony-informative site | Singleton |
| --- | --- | --- | --- | --- | --- |
| **matK** | 49 | 659 | 66 | 27 | 39 |
| **rbcL** | 30 | 1184 | 24 | 6 | 18 |

Similarly, 24 variable sites, 6 parsimony-informative sites, and 18 singleton sites were recorded in polymorphism site analysis of rbcL. The highly conserved region was found (Region: 1024-1196) in rbcL sequence alignment (CAAGATTGGGTTTCTATGCNAGGNGTTATTCCCGNGGCTTCAGGNGGTATTCATGTTTGGCANANGCCTGCCCTAACTGAAATCTTTGGAGATGATTCCGTACTACAGTTCGGTGGAGGAACTTTAGGACACCCTTGGGGAAATGCACCCGGTGCGGTAGCNAATCGGGT AGC). The 18 singleton variable sites were located at position 17, 183, 220, 228, 246, 291, 387, 430, 456, 474, 600, 624, 657, 695, 780, 792, 909, 1023, and 6 parsimony informative sites at position 177, 195, 273, 531, 729, 1197 (Table 2).

This study evaluates the substitution of different bases in analyzed regions on entire codon positions (1st+ 2nd + 3rd nucleotide). In general, transitional substitution is higher than transversional substitution in both rbcL and matK regions. The candidate barcode gene, rbcL, showed a higher transitional substitution rate than matK. The highest substitution rate from A to G and T to C was noted in the rbcL region (14.62%). Similarly, the changing frequency from G to A and C to T of rbcL (18.91%) was found higher as compared to the matK (16.77%) region (Table 3).

**Table 3.** The maximum likelihood estimate of the substitution matrix in the constructed phylogenetic tree of the genus *Aloe*.

| Gene | matK |  |  |  | rbcL |  |  |  |
| --- | --- | --- | --- | --- | --- | --- | --- | --- |
| Nucleotide | A | T/U | C | G | A | T/U | C | G |
| **A** | - | 8.87 | 3.98 | **7.53** | - | 4.64 | 3.59 | **14.62** |
| **T/U** | 8.87 | - | **7.53** | 3.98 | 4.64 | - | **14.62** | 3.59 |
| **C** | 8.87 | **16.77** | - | 3.98 | 4.64 | **18.91** | - | 3.59 |
| **G** | **16.77** | 8.87 | 3.98 | - | **18.91** | 4.64 | 3.59 | - |

**NOTE-** Each entry is the probability of substitution (*r*) from one base (row) to another base (column). Substitution patterns and rates were estimated under the Tamura (1992) model (**Tamura, 1992**). Rates of different transitional substitutions are shown in **bold**, and those of transversions substitutions are shown in *italics*. Evolutionary analyses were conducted in MEGA11.

### Genetic diversity

Genetic diversity is the variation at the nucleotide level, including variations across individuals and populations within each species. The significance of genetic diversity was verified by nucleotide diversity and neutrality tests; Tajima’s D, Fu & Li’s D, and Fu & Li’s F test **(Jiang et al., 2022)**. According to the observed cumulative Eta value, the matK sequences exhibited the highest level of genetic variety, with 67 mutations found in the matK sequence, compared to the rbcL sequence (Eta value = 24). Moreover, the nucleotide and haplotype diversity of matK was higher than that of rbcL. Tajima’s D, Fu & Li’s D, and Fu & Li’s F test statistics were statistically negative but not significant (P > 0.02). The results suggested these populations were stable, with no recent bottleneck or rapid population expansion **(Cao et al., 2019)**.

All the results showed significant genetic variation in *Aloe* species for candidate nucleotide sequences. DNA sequences were evaluated for population size changes to observe nucleotide mismatch distribution across different *Aloe* species, which enriched the findings of genetic diversity among species (Table 4; Figure 1).

**Table 4.**
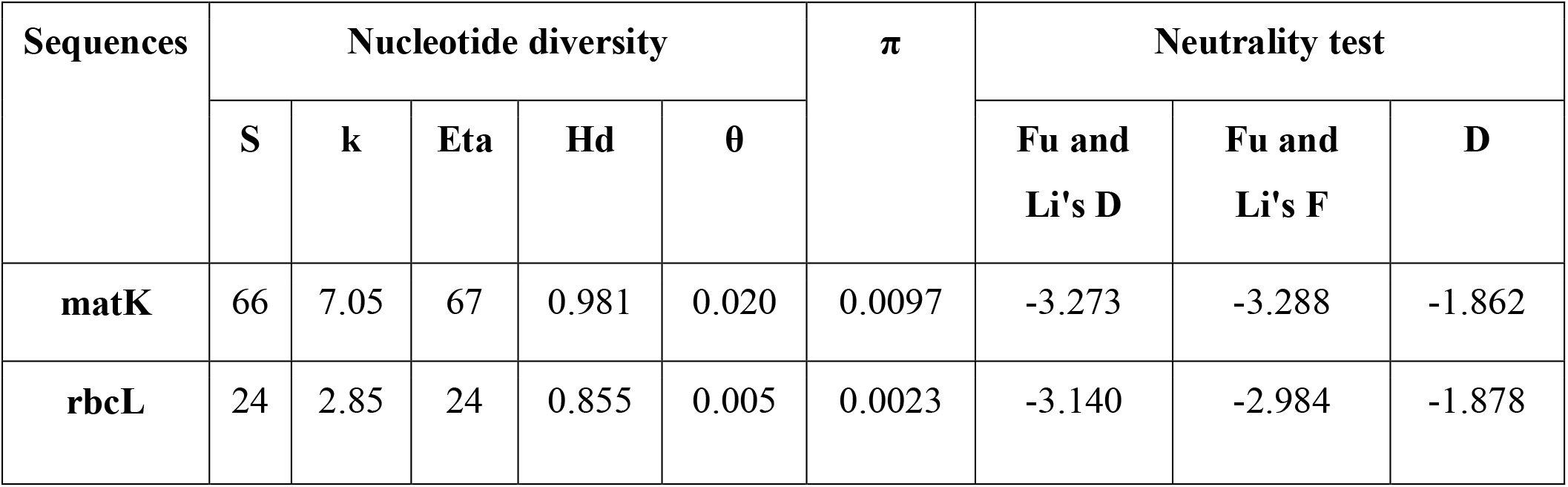
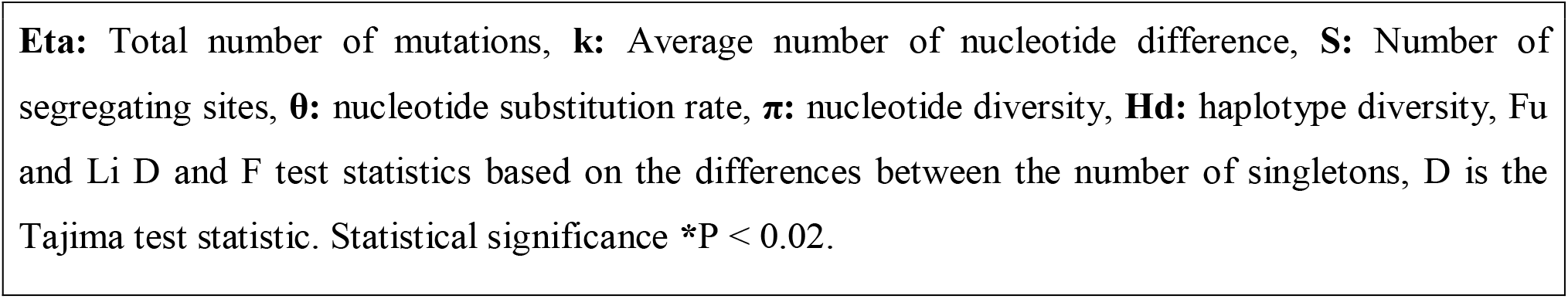
Genetic diversity calculation of *Aloe* plants based on candidate barcode sequences.

| Sequences | Nucleotide diversity | | | | | $\pi$ | Neutrality test | | |
| --- | --- | --- | --- | --- | --- | --- | --- | --- | --- |
| | S | k | Eta | Hd | $\theta$ | | Fu and Li's D | Fu and Li's F | D |
| **matK** | 66 | 7.05 | 67 | 0.981 | 0.020 | 0.0097 | -3.273 | -3.288 | -1.862 |
| **rbcL** | 24 | 2.85 | 24 | 0.855 | 0.005 | 0.0023 | -3.140 | -2.984 | -1.878 |

**Eta:** Total number of mutations, **k:** Average number of nucleotide difference, **S:** Number of segregating sites, **θ:** nucleotide substitution rate, **π:** nucleotide diversity, **Hd:** haplotype diversity, Fu and Li D and F test statistics based on the differences between the number of singletons, D is the Tajima test statistic. Statistical significance \*P < 0.02.

**Figure 1.**
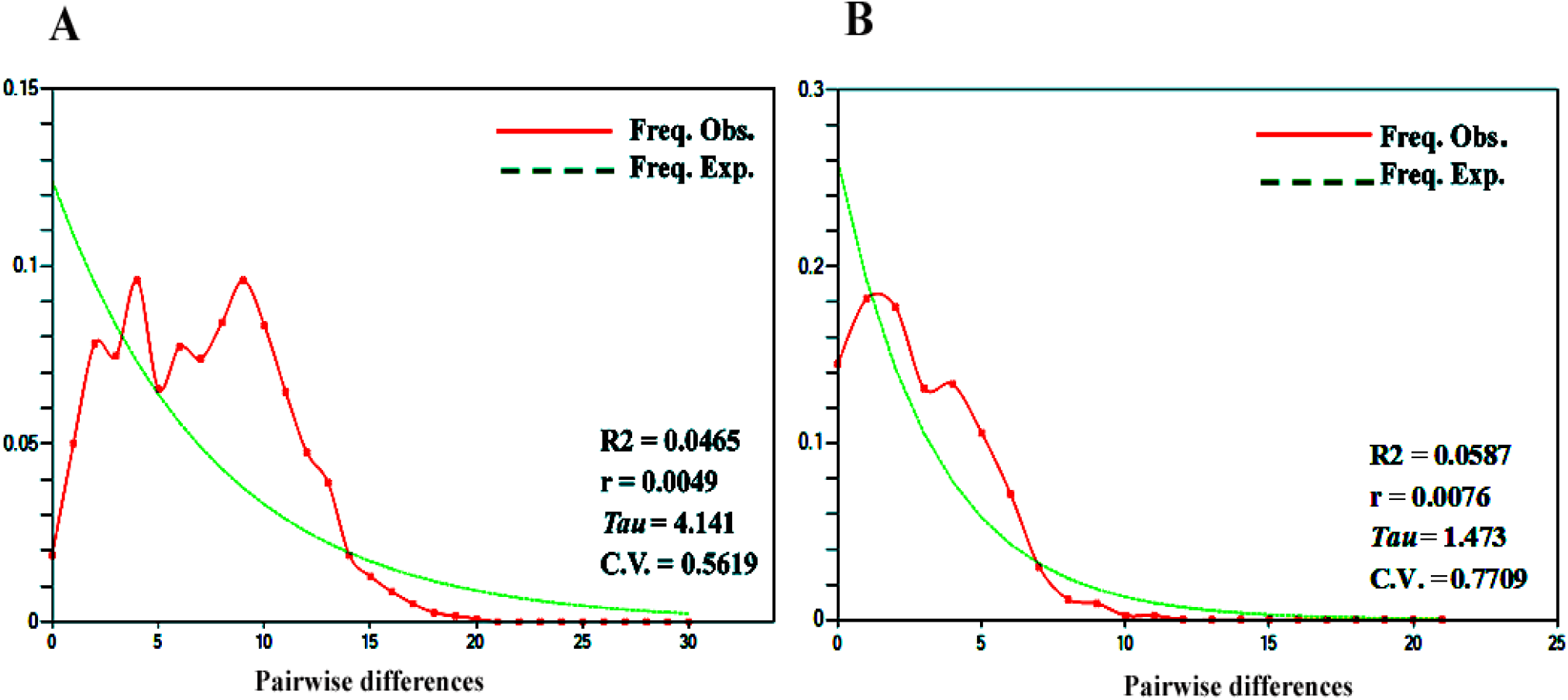
Mismatch distribution shows the pairwise differences frequency based on matK (A) and rbcL (B) sequences. The X-axis shows the pairwise differences, and the Y-axis shows the frequency. R2 Ramos-Onsins and Rozas statistics; r Raggedness statistic; Tau Date of the Growth or Decline measured of mutational time; CV Coefficient of variation.

### Phylogenetic Analysis

This study used MEGA11 software to identify the evolutionary relationship based on matK and rbcL genes using the Maximum Parsimony method (Figure 2). The variability among rbcL and matK regions is from 0.00 to 0.01 and 0.03, respectively. The 49 *Aloe* samples were allocated into four major clades based on matK sequence: Clade I is the largest and represented by *Aloe vera;* Clade II is represented by *Aloe boylei, Aloe suprafoliata, Aloe welwitschii*, and *Aloe verecunda*; Clade III is represented by *Aloe ciliaris*; and Clade IV is represented by *Aloe ferox* and *Aloe arborescens* (Figure 3). Similarly, the 29 *Aloe* samples were distributed into two major clades based on rbcL sequence: Clade I is represented by *Aloe vera*; Two *Aloe* species, namely, *Aloe arborescens* and *Aloe maculate* represent Clade II (Figure 4). Considering the topological structure of the evolutionary tree, both chloroplast genes (matK and rbcL) exhibit better identification ability at the species level **(Grace et al., 2015; Han, 2016; Alaklabi et al., 2021; Algarni et al., 2022a; b)**.

**Figure 2.**
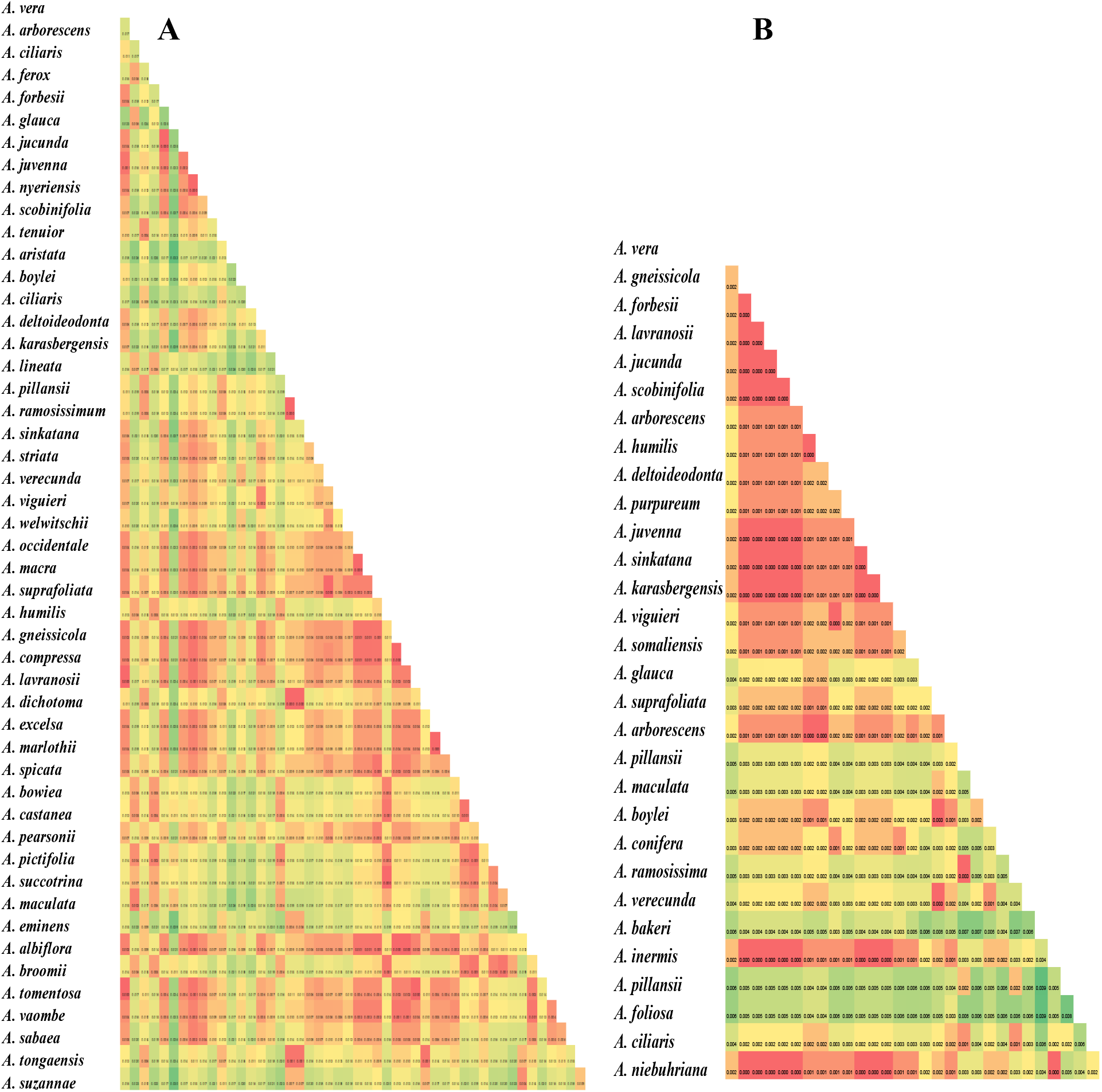
Pairwise genetic distance heatmap based on candidate barcode sequences (matK (A) and rbcL (B). Analysis was conducted using the Tamura 3-parameter model (Tamura, 1992).

**Figure 3.**
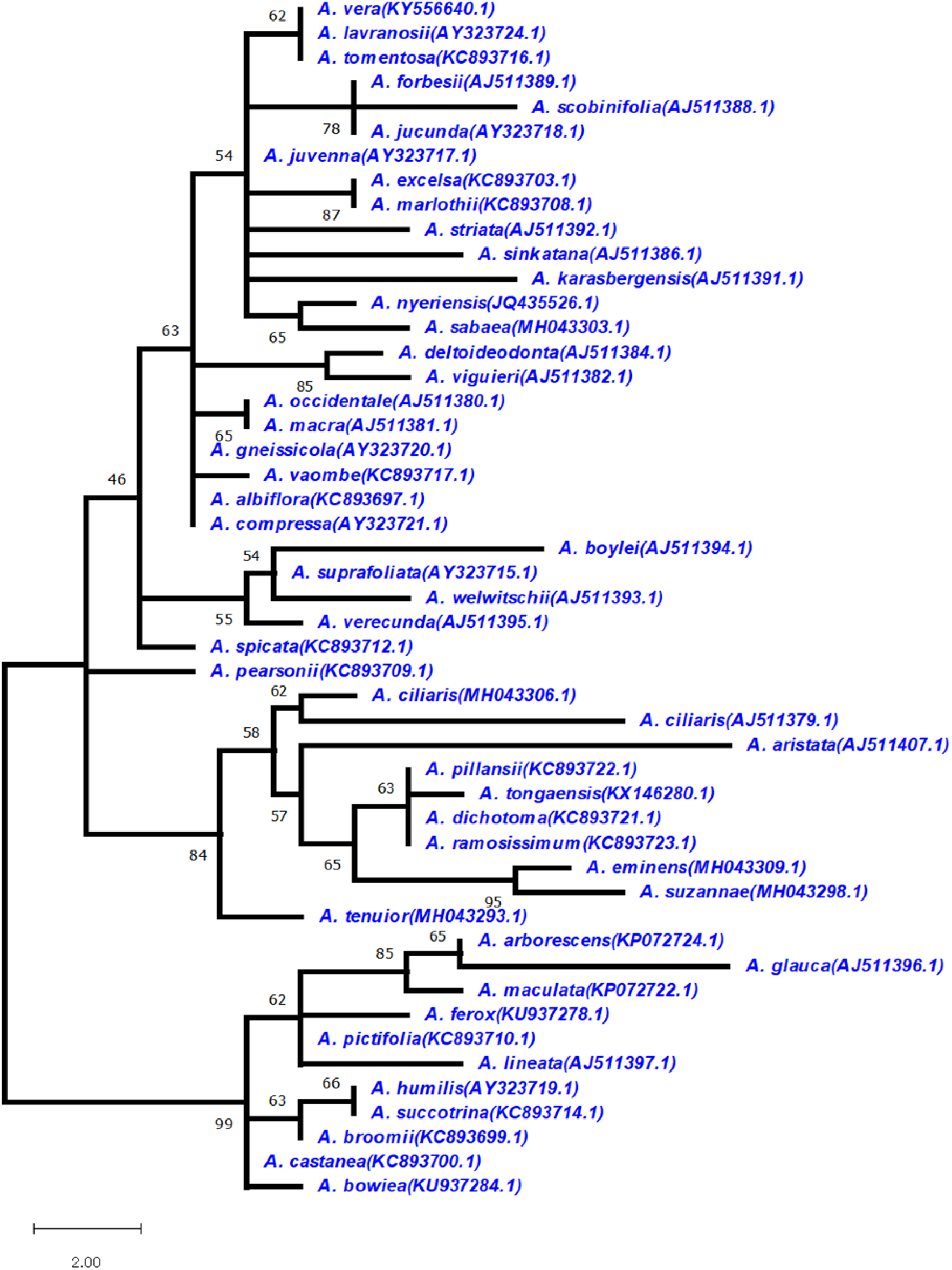
The evolutionary history was inferred using the Maximum Parsimony method. This analysis involved 49 nucleotide sequences of the matK gene and a total of 835 positions in the final dataset. The Numbers on the branches represent more than or equal to 40 percent support after the 10,000 bootstrap replications test.

**Figure 4.**
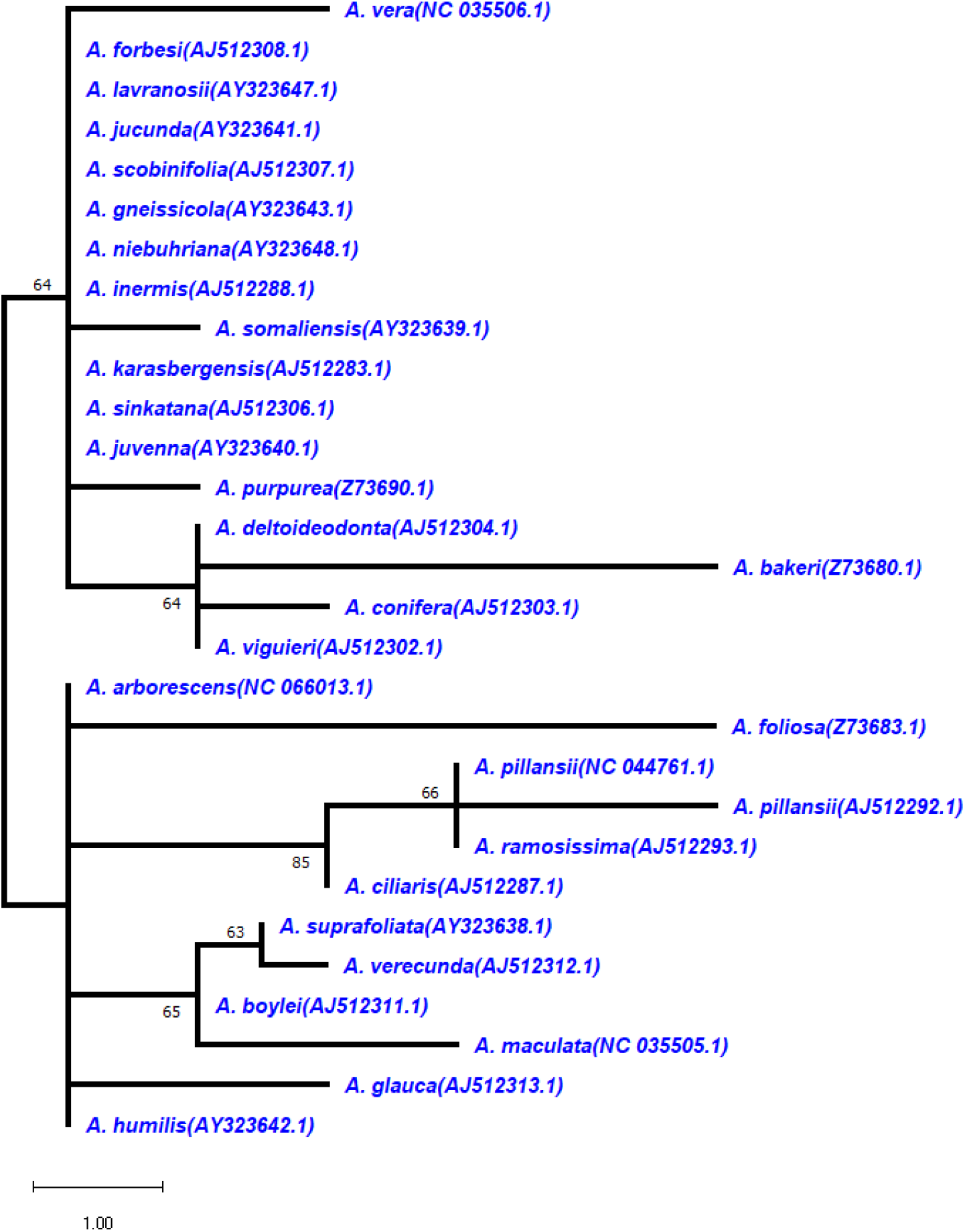
The evolutionary history was inferred using the Maximum Parsimony method. This analysis involved 29 nucleotide sequences of the rbcL gene and a total of 1265 positions in the final dataset. The Numbers on the branches represent more than or equal to 40 percent support after the 10, 000 bootstrap replications test.

### Analysis of barcoding gap

An ideal DNA barcoding sequence for species identification should satisfy that inter-specific genetic variation is significantly higher than intra-specific genetic variation **(Li et al., 2021)**. The individual chloroplast gene sequences were evaluated to verify the applicability of candidate sequences, and the barcoding gap was analyzed according to the frequency distribution histogram based on pairwise distance divergence (Figure 5). The maximum K2P intra-specific genetic distances were much smaller than the minimum inter-specific genetic distance of all samples, and due to some overlaps of intra- and inter-specific variation, no noticeable barcoding gap was observed in matK and rbcL sequences **(Han, 2016)**.

**Figure 5.**
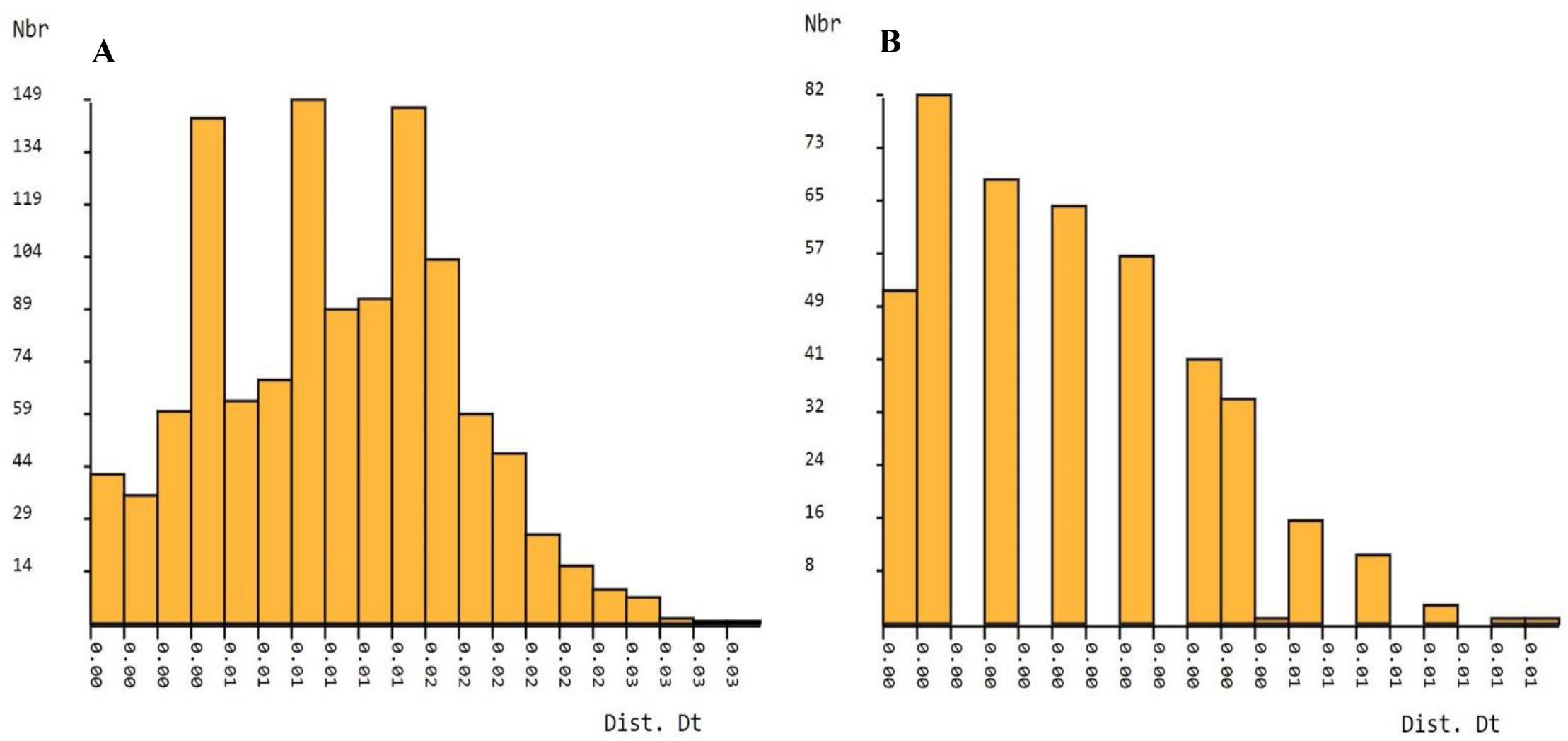
Histogram showing pairwise distance divergence (%) generated by Automatic Barcode Gap Discovery (ABGD) based on matK (A) and rbcL (B) sequences of *Aloe species*. The X-axis shows the genetic distance distribution, and the Y-axis shows Nbgroups. Nbgroups is the number of species as identified by ASAP in the corresponding partition.

### Specific barcodes based on SNP sites

Based on SNP sites, species-specific barcodes were developed, and the appropriate fragments were blasted into the NCBI database **(Li et al., 2021)**. Based on the rbcL sequence, the specific barcode of *Aloe vera* was generated. Information about particular barcodes of four *Aloe* species; *Aloe boylei, Aloe ciliaris, Aloe karasbergensis*, and *Aloe sinkatana* was obtained based on the matK sequence. Electronic equipment can scan two-Dimensional code from DNA fragments that can be used for species identification. It can provide theoretical support for future research. The species-specific barcode was generated using the Two-Dimensional code coding method, which converted a unique barcode sequence into a two-dimensional barcode image, which helped convert barcode information (Figure 6).

**Figure 6.**
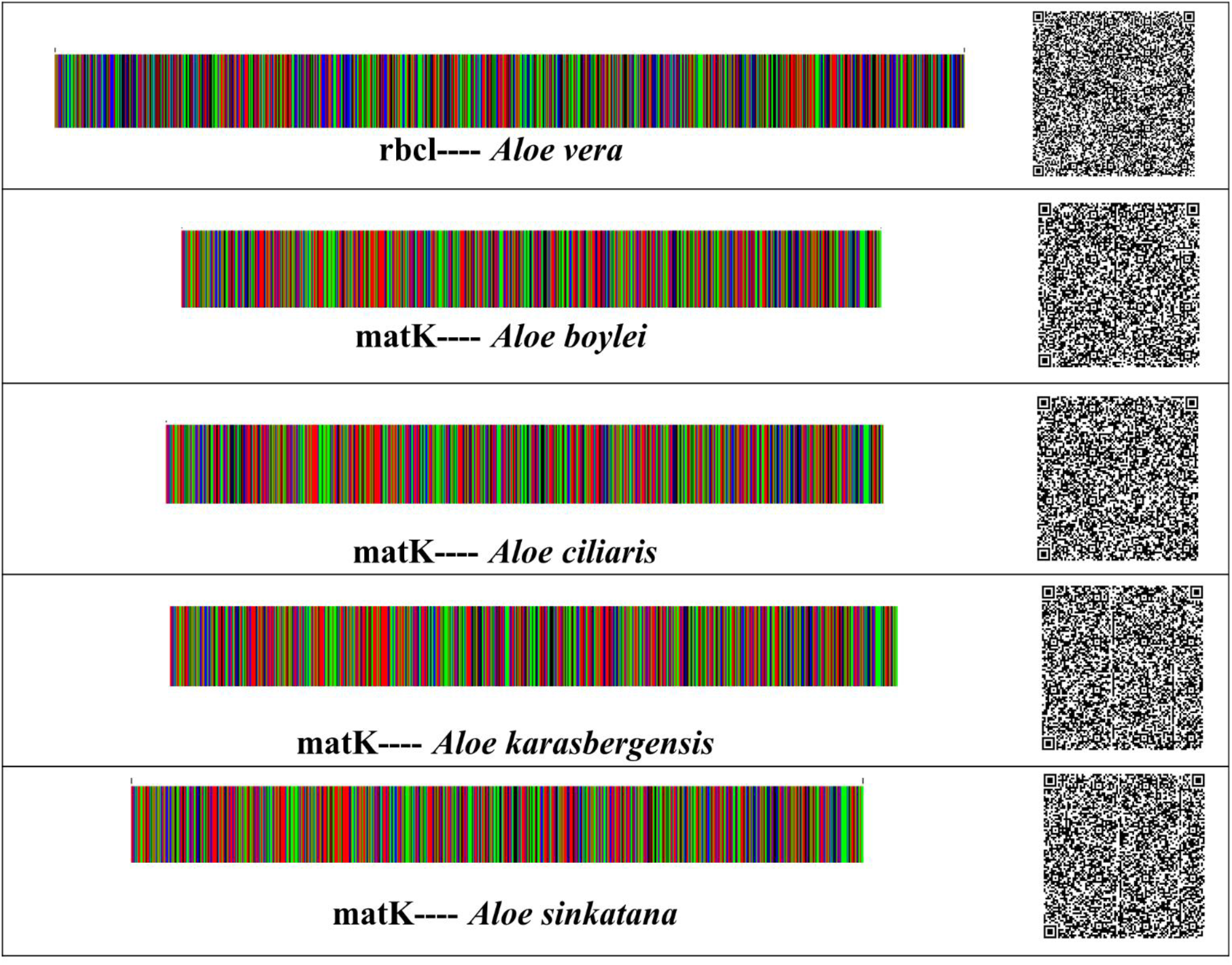
DNA barcodes and two-dimensional DNA barcodes of *Aloe sp*. based on matK and rbcL genes. Base A in green, base T in red, base C in blue, and base G in black.

## CONCLUSIONS

DNA barcode can be utilized for species identification by amplifying DNA fragments shared by all species. These advanced techniques allow the identification and authentication of plant species, plant-based raw materials, and their traceability in the final products. These complementary techniques enable dealing with adulterations encountered mainly in the cosmetic, nutraceutical, and food industries to guarantee consumer transparency. The CBOL recommends matK and rbcL as universal barcodes in the plant kingdom. The findings of sequence studies on average GC content showed that the GC content of *Aloe* sp. candidate barcode sequences was less than AT content. The phylogenetic analyses showed that the identification ability of matK and rbcL was average on the *Aloe* genus level. The limited availability of sequences from some species could affect the precision of the study. Additional *in silico* study should be performed with a higher sequence number to raise the discrimination ability of DNA barcode loci. This study extends the application of the matK and rbcL sequences in DNA barcoding of the genus *Aloe*. Based on the SNP analysis, the species-level “specific DNA barcodes” of *Aloe* were successfully developed. Also, specific DNA barcodes provide a good solution for species discrimination of a given taxonomic group based on DNA sequence data establishing the base for the conservation, evaluation, and utilization of *Aloe* germplasm.

